# EmbedGEM: A framework to evaluate the utility of embeddings for genetic discovery

**DOI:** 10.1101/2023.11.24.568344

**Authors:** Sumit Mukherjee, Zachary R McCaw, Jingwen Pei, Anna Merkoulovitch, Tom Soare, Raghav Tandon, David Amar, Hari Somineni, Christoph Klein, Santhosh Satapati, David Lloyd, Christopher Probert, Insitro Research Team, Daphne Koller, Colm O’Dushlaine, Theofanis Karaletsos

**Affiliations:** Insitro inc., California, USA; Center for Machine Learning, Georgia Institute of Technology, Georgia, USA; Chan-Zuckerberg Insitiative, California, USA

**Keywords:** GWAS, Representation learning, Multivariate traits

## Abstract

Machine learning (ML)-derived embeddings are a compressed representation of high content data modalities. Embeddings can capture detailed information about disease states and have been qualitatively shown to be useful in genetic discovery. Despite their promise, embeddings have a major limitation: it is unclear if genetic variants associated with embeddings are relevant to the disease or trait of interest. In this work we describe EmbedGEM (**Embed**ding **G**enetic **E**valuation **M**ethods), a framework to systematically evaluate the utility of embeddings in genetic discovery. EmbedGEM focuses on comparing embeddings along two axes: heritability and disease relevance. As measures of heritability, we consider the number of genome-wide significant associations and the mean *χ*^2^ statistic at significant loci. For disease relevance, we compute polygenic risk scores for each embedding principal component, then evaluate their association with high-confidence disease or trait labels in a held-out evaluation patient set. While our development of EmbedGEM is motivated by embeddings, the approach is generally applicable to multivariate traits, and can readily be extended to accommodate additional metrics along the evaluation axes. We demonstrate EmbedGEM’s utility by evaluating embeddings and multivariate traits in two separate datasets: i) a synthetic dataset simulated to demonstrate the ability of the framework to correctly rank traits based on their heritability and disease relevance, and ii) a real data from the UK Biobank including metabolic and liver-related traits. Importantly, we show that greater disease relevance does not automatically follow from greater heritability.

## Introduction

Representation learning is a crucial aspect of modern day machine learning (ML), whose aim is to discover compact and informative representations of high-dimensional data. ML-derived representations are often simply called *embeddings*. Deep neural networks have emerged as a powerful tool for representation learning, demonstrating remarkable success across various domains. Representation learning was first introduced in the form of unsupervised pre-training [12], where a deep neural network was trained on unlabeled data to initialize weights for subsequent supervised learning tasks. Autoencoders inaugurated the next generation of representation learning algorithms, which are neural networks trained to reconstruct their input, enabling the learning of compact and informative representations [21]. More recently, self-supervised learning has gained attention as a promising approach, where the model learns representations by maximizing agreement between differently augmented views of the same data [6, 5]. Self-supervised pre-training has significantly improved performance on tasks including image classification, natural language processing, and recommendation systems [9].

In recent years, embeddings have become increasingly used in genetic discovery [17, 14, 7, 23, 18, 25], identifying novel associations between genetic variants and disease indications. While some studies have heuristically evaluated the utility of embeddings for genetic discovery, there is currently little systematic work evaluating their added value. In [25], genome-wide association studies (GWASs) were performed on relatively uncorrelated embeddings extracted from *β* variational autoencoders [11]. The authors established disease relevance by generating polygenic risk scores (PRSs) from embedding-associated variants, and demonstrating improved discrimination between cases and controls in independent cohorts as compared with PRSs composed of variants associated with existing multivariate traits. While this was not developed into an explicit framework for evaluating the disease relevance of embeddings, it served as a motivation for our approach. In [14], the authors conducted univariate GWAS of embedding principal components (PCs). Subjects with extreme values in PC space were inspected to qualitatively interpret the PCs, and related phenotypes were identified by performing PheWAS [8] on the PCs. Similarly, [18, 23] performed GWAS on each dimension of the embeddings and conducted genetic correlation analysis with other relevant traits to evaluate the relevance of the different embedding dimensions. Mukherjee and colleagues generated unsupervised 2D representations learned from bulk RNA-Seq data and used a metric based on patient distance from the prototypical baseline as a quantitative phenotype in GWAS [17]. The authors validate the metric against other known clinical metrics to quantify disease severity.

Unlike prior works which focused on simply performing genetic discovery with embeddings, here we propose a formal framework for evaluating the utility of embeddings for genetic discovery. EmbedGEM evaluates embeddings along two dimensions: heritability and disease relevance. Using both simulated and real datasets, we demonstrate the utility of EmbedGEM for evaluating embeddings and multivariate traits more broadly. Finally, we release a software implementation of EmbedGEM (https://github.com/insitro/EmbedGEM) along with a tutorial on how to use it.

## Preliminaries

### Mathematical formulation of representation learning

Let *X* denote the input data space and *Y* denote a lower-dimensional learned representation space. Given a dataset of the form 𝔻 = (*x*_1_, *l*_1_), (*x*_2_, *l*_2_), …, (*x*_*n*_, *l*_*n*_), where *x*_*i*_ ∈ *X* is an input sample (e.g. image) and *l*_*i*_ is an optional corresponding label, the goal of representation learning is to find a mapping *f* : *X* → *Y* that captures meaningful features and the underlying structure of the data. This is often formulated as a supervised learning problem, where the objective is to minimize a loss function (*L g* ∘ *f* (*x*_*i*_), *l*_*i*)_ that measures the discrepancy between the predicted labels using the learned representations *g* ∘ *f* (*x*_*i*_) and the true label *l*_*i*_. Here *g*(·) is a projection that maps the representation *f* (*x*_*i*_) into the space where the loss is calculated. In the absence of meaningful labels, representation learning can also be formulated as an unsupervised or self-supervised learning problem where the loss function is of the form *L(g* ∘*f* (*x*_*i*_), *x* (e.g. for an auto-encoder), or *L (g* ∘ *f* (*x*_*i*1_), *g* ∘ *f* (*x*_*i*2_), where *x*_*i*1_ and *x*_*i*2_ are two different views of *x*_*i*_ (e.g. for SimCLR). In any case, the primary output of the representation learning algorithm is *y*_*i*_ = *f* (*x*_*i*_), which is a lower dimension representation of the input (commonly referred to as embeddings).

### Genome-wide association studies (GWASs)

GWAS [22] involves analyzing a large number of single nucleotide polymorphisms (SNPs) across the genome to identify associations between specific genetic variants and phenotypes of interest. Mathematically, GWAS of a quantitative trait is posed as a linear regression problem, where the phenotype *Y* is regressed on the genotype *G*, i.e. the number of risk alleles an individual carries at a genomic location, adjusting for a vector of covariates *X*:

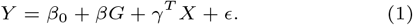

Here *β* is the regression coefficient of interest, *γ* is a vector of coefficients for the adjustment variables, and *ϵ* a residual with mean zero and finite variance. Variants associated with the phenotype are identified by rejecting the null hypothesis *H*_0_ : *β* = 0. Due to the large number of variants in the genome, standard practice is to run a separate association test for each SNP *G*.

### LD-based Clumping

Nearby genetic variants on a chromosome tend to be inherited together, leading to correlations among variants known as Linkage Disequilibrium (LD). Clumping is the process by which variants in high LD with the most significant variant in a region, the *index variant*, are pruned, or removed, to reduce redundancy [1]. Clumping can be performed using common statistical genetic software, notably `plink` [19]. Mathematically, clumping can be formulated as a greedy selection process, where for each LD region ℛ_*i*_, only the variant with the lowest p-value is retained:

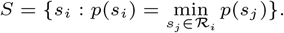

Here *S* is the clumped set of variants, *p*(*s*_*i*_) is the p-value of variant *s*_*i*_, and ℛ _*i*_ is the set of variants in the same LD region as *s*_*i*_. The LD neighborhood of a variant is typically defined by the set of variants correlated at an *R*^2^ threshold of 0.5 or 0.1.

### Polygenic Risk Scores (PRS)

Polygenic risk scores (PRS) [16] have emerged as a powerful tool in genetic epidemiology for predicting an individual’s risk of developing complex traits and diseases. A PRS is calculated by summing the contributions of multiple genetic variants across the genome, weighted by their association with the phenotype of interest from (1)

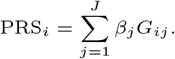

Here PRS_*i*_ is the polygenic score for subject *i, G*_*ij*_ represents the number of risk alleles subject *i* carries at genetic variant *j*, and *β*_*j*_ represents the estimated genetic effect size. By aggregating the effects of multiple genetic variants, PRS provides a quantitative measure of genetic predisposition to a particular trait or disease.

## Materials and methods

The EmbedGEM workflow, consisting of heritability and disease relevance evaluations is summarized Figure 1. The heritability evaluation comprises of deriving multivariate GWAS summary statistics from univariate GWAS summary statistics of the orthogonalized embedding PCs (section 3.1.1). These summary statistics are used to compute metrics that quantify the total heritability of the embeddings (section 3.1.2). For evaluating disease relevance, we assess the performance of polygenic risk scores, composed of variants associated with the orthogonalized embedding dimensions, for predicting a disease trait of interest (section 3.1.3).

**Fig. 1.**
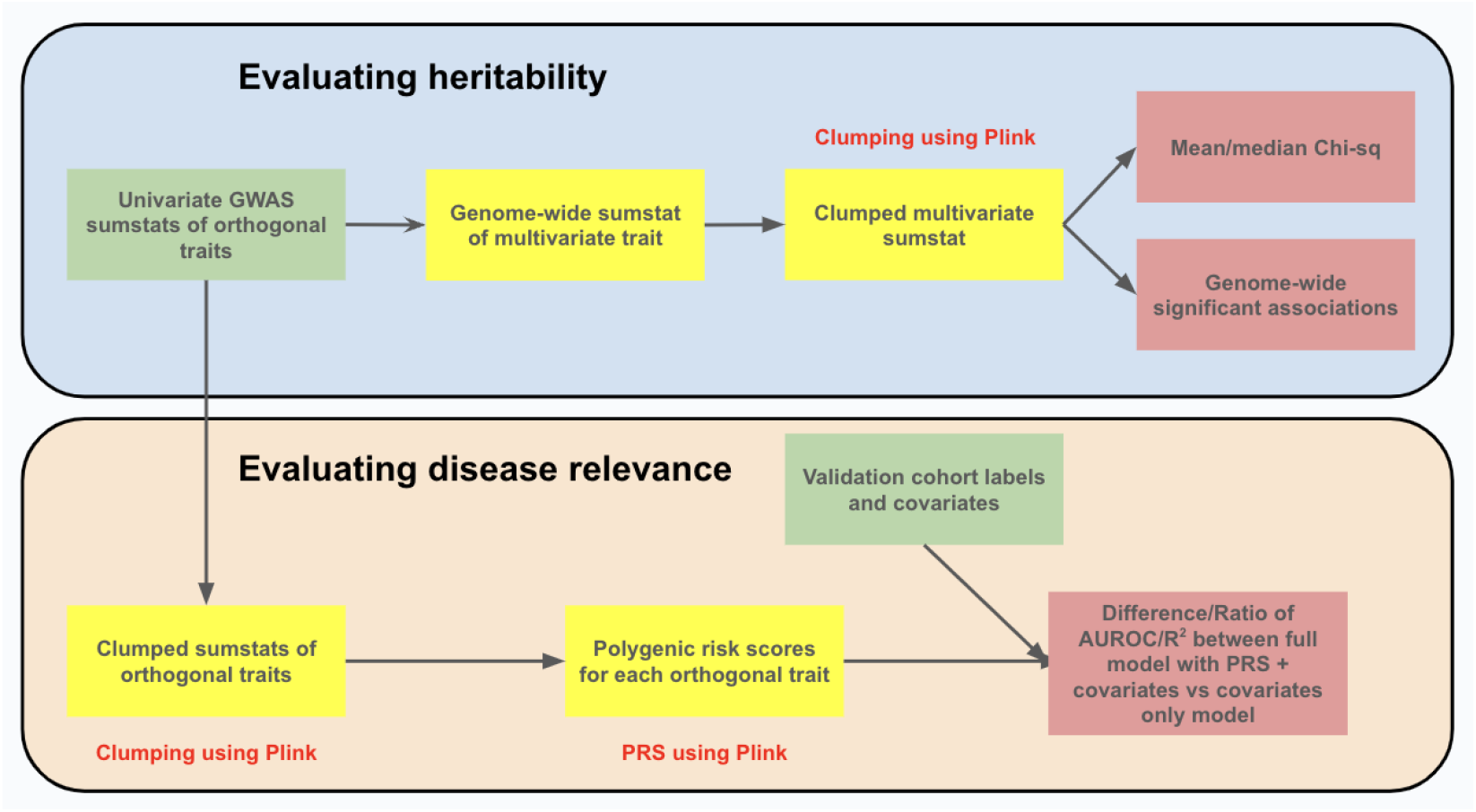
Overview of EmbedGEM’s genetic validation workflow. Green boxes indicate inputs, yellow boxes indicate intermediates, and red boxes indicate final output metrics. Parts of the workflow that utilize `plink` commands are highlighted using red text.

Aside from introducing the different components of the workflow, this section introduces the datasets used to evaluate EmbedGEM. First we describe a simulated dataset that we use to demonstrate the ability of EmbedGEM to correctly order different embeddings (section 3.2). Then we introduce a real world example, along with the methods used to process the data and generate embeddings (section 3.3).

### Evaluation methods

#### GWAS methodology for multivariate traits

Given *K* traits or embedding dimensions (*Y*_1_, …, *Y*_*K*_), we perform Principal Component Analysis (PCA) to obtain *K* orthogonal principal components (PCs), denoted as PC_1_, …, PC_*K*_. Subsequently, we conduct single-trait GWAS for the first *m* ≤ *K* PCs (where *m* was either user-selected or determined through an automated procedure) using the following association model:

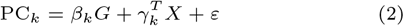

Here PC_*k*_ is *k*th PC, *G* is genotype at the variant of interest, and *X* is a vector of covariates, such as age, sex, and ancestry PCs. The effect of genotype on the *k*th PC is captured by *β*_*k*_. Because the principal components are orthogonal by construction, the per-component Wald statistics 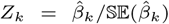 are independent and multivariate normal asymptotically (i.e. as the number of subjects → ∞). Under the null hypothesis of no association, the combined test statistic 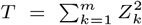 follows a central 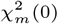 distribution with *m* degrees of freedom [2]. We use 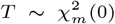 to evaluate the hypothesis:

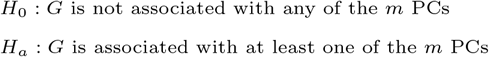

Note that although the embeddings were orthogonalized via PCA in this work, EmbedGEM does not depend on any particular method of orthogonalizing the traits.

#### Evaluating heritability

The goal of evaluating heritability is to compare the different multivariate traits (e.g. embeddings) in terms of their ability to manifest genetic associations. In EmbedGEM, we examine the following summary statistic-based metrics, which are commonly reported in the field:

- Number of independent genome-wide significant (GWS) variants (p-value ≤ 5 × 10^−8^) after LD-based clumping.
- The mean and median *χ*^2^ statistics of the independent

(clumped) GWS variants.

The mean *χ*^2^ statistic at independent GWS variants is an assessment of signal strength. An approach that provides a higher mean *χ*^2^ provides better power, allowing for detection of a genetic association with a smaller sample size. The total number of GWS variants instead gauges the heritability of the trait. We provide a theoretical rationale for using these metrics in Section 6 of the **Supplementary Methods**. We also provide empirical experiments to demonstrate their relationship to the commonly used LD-score regression (LDSC) method for evaluating heritability [3].

#### Evaluating disease relevance

The purpose of the disease relevance evaluation is to assess the extent to which the embedding associated variants are predictive of an outcome of interest. More specifically, we evaluate the strength of association between the orthogonalized trait PRSs and user-provided disease labels by comparing a full disease prediction model, which includes all PRSs:

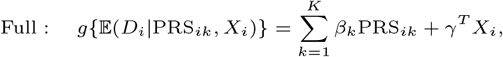

with a reduced model, which excludes them:

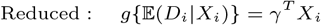

Each equation represents a generalized linear model (GLM) in which *D*_*i*_ is the disease or evaluation trait for the *i*th subject, (PRS_*ik*_) that subject’s PRS with respect to the *k*th embedding, and *X*_*i*_ a vector of covariates, which can differ from those included in the GWAS model (2). For a binary trait, logistic models are fit, while for a continuous trait, linear regression models are fit.

From a fitted GLM, a prediction 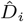of the disease trait is obtained, which can be compared with the observed *D*_*i*_ using various metrics. For binary traits, we calculate the area under the receiver operating characteristic (AUROC) and the area under the precision-recall curve (AUPRC). For continuous traits, we calculate the square correlation and the mean absolute prediction error (MAE). Metrics for the full and reduced models are compared with respect to a contrast Δ (e.g. difference or ratio). To assess whether addition of the PRSs significantly improves disease relevance, we estimate the distribution of Δ under the null by first permuting the PRSs, rendering them non-informative, then taking *B* bootstrap resamples of the data (with replacement). For each resample *b*, the full and reduced models are fit, and metric contrast Δ_*b*_ calculated. Letting Δ_obs_ denote the observed contrast (on the unpermuted data), the final p-value is:

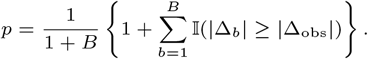

#### Simulated dataset

To validate that the proposed workflow can disentangle heritability and disease relevance, we simulated orthogonalized embeddings and outcome phenotypes in a setting where the generative architecture is known. We sampled variants in linkage equilibrium at *R*^2^ ≤ 0.1 for ∼350K unrelated subjects of white British ancestry from the UK Biobank. Each PC was generated from an infinitesimal model:

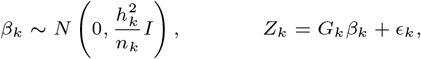

where *G*_*k*_ is the set of genetic variants affecting *Z*_*k*_, standardized to have mean 0 and variance 1, *β*_*k*_ is the effect size vector, 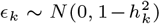 is a residual, 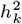 is the heritability of *Z*_*k*_, and *n*_*k*_ is the number of causal variants. Each simulated embedding depended on *n*_*k*_ = 1000 variants, and effect sizes for *Z*_1_ and *Z*_2_ were drawn independently, such that the correlation of *Z*_1_ and *Z*_2_ had expectation zero.

We simulated a continuous disease liability trait *Y* from the following model:

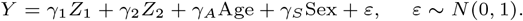

The liability *Y* was simulated such that each PC, age, and sex independently explained 10% of the variation. Three scenarios were considered:

i. **High heritability and high disease relevance**: Here each embedding PC had 20% heritability, and 20% of the variation in *Y* was explained by *Z*_1_ and *Z*_2_.
ii. **High heritability but low disease relevance**: Here *Y* was permuted such that *Z*_1_ and *Z*_2_ remained heritable but the outcomes no longer depended on the embedding PCs.
iii. **Low heritability and low disease relevance**: Here all of *Z*_1_, *Z*_2_, and *Y* were permuted such that neither the embedding PCs nor the outcomes were heritable.

### Real-world dataset

#### Non-Alcoholic Fatty Liver Disease (NAFLD)

NAFLD is a prevalent and complex chronic liver condition, characterized by the accumulation of excess fat in the liver of individuals who consume little to no alcohol [24]. It is considered the hepatic manifestation of metabolic syndrome. NAFLD encompasses a wide spectrum of liver damage, ranging from simple steatosis to non-alcoholic steatohepatitis (NASH), advanced fibrosis, and cirrhosis. The global prevalence of NAFLD is estimated to be approximately 25%, making it a significant public health concern.

The UK Biobank (UKB) is a large-scale biomedical database and research resource, containing in-depth genetic and health information from 500K UK individuals aged 40-69 years [4]. The resource includes data on a wide range of health-related outcomes, which are being further enriched via linkage to diverse medical, social, and environmental records. Neck-to-knee MRIs from UKB have previously been used to extract various adiposity and organ traits [15, 20]. We followed the method outlined in [15] to process the neck-to-knee MRIs and impute adiposity traits for approximately 36K patients. These deep imputation models were also the source for our supervised embeddings.

In consultation with subject-matter experts, we identified 203 fields that have a known or suggested association with the disease of interest (included in supplementary materials). These traits were grouped into: i) nuclear magnetic resonance metabolomics, ii) abdominal composition variables, iii) blood biochemistry markers. We also identified 48 covariates that might affect traits of interest independently from the disease processes of interest. The list of covariates were grouped into: i) body size, ii) addiction information, iii) medical status information, iv) medication usage information, v) baseline characteristics.

To curate a cohort for evaluating disease relevance, we identified 1774 NAFLD cases and 2209 NAFLD controls using ICD-10 codes found in UKB. These individuals were removed from the discovery cohort (individuals with neck-to-knee MRIs).

#### Learning supervised embeddings

Supervised embeddings were learned using the process described in [15] for each adiposity trait separately (see list of traits in appendix). The modeling procedure involved replacing the last layer of an ImageNet pre-trained ResNet-50 [10] model with a linear layer (to obtain regression predictions). The model was then fine-tuned end-to-end using 80% of the labeled data for training and 20% of the data for testing. An adaptive learning rate schedule was used for the Adam optimizer, as stated in [15]. 2048 dimensional embeddings were extracted from the flattened output of the last convolutional layer.

#### Type 2 Diabetes

In Section 4 of the Supplementary Methods, we present an analysis on the relevance of embeddings extracted from color fundus images to type 2 diabetes status in the UKB.

## Results

### EmbedGEM correctly distinguishes embedding heritability from disease relevance

As described in Section 3.2, we simulated data from three genetic architectures in order to demonstrate the utility of EmbedGEM namely, i) high heritability and high disease relevance, ii) high heritability and low disease relevance, iii) low heritability and low disease relevance. Table 1 demonstrates that EmbedGEM correctly differentiates between heritability and disease relevance. For instance, in both of the high heritability scenarios, we observe a substantial number of genome-wide significant (GWS) associations and a significantly elevated mean *χ*^2^. However, only in the case of the high disease relevance trait do the embedding PRSs significantly improve association with the disease liability, as evidenced by the significant *r*^2^ and MAE. Note that the magnitudes of the *r*^2^ and MAE should be interpreted with caution since the baselines for different architectures can, and in this case do, differ.

**Table 1.** EmbedGEM correctly distinguishes embedding heritability from disease relevance. The table shows the results of our EmbedGEM for three scenarios, differing by the heritability of the embeddings and of the final disease labels. The framework correctly orders the three scenarios in terms of heritability and disease relevance. The scenarios with high embedding heritability have a high mean *χ*^2^ & number of genome-wide significant (GWS) hits, while the scenario with low heritability has no GWS hits. The disease relevance is also correctly identified, with only the high disease relevance trait exhibiting a significant *r*^2^ and MAE.

| Genetic Architecture |  | Disease relevance |  |  |  | Heritability |  |
| --- | --- | --- | --- | --- | --- | --- | --- |
| Heritability | Disease Relevance | $r^2$ ratio | $r^2$ p-value | MAE ratio | MAE p-value | No. of hits | Mean $\chi^2$ |
| High ↑ | High ↑ | 1.20 | 0.001 | 0.97 | 0.001 | 2207 | 100.29 |
| High ↑ | Low ↓ | 3.25 | 0.434 | 0.999 | 0.513 | 2207 | 100.29 |
| Low ↓ | Low ↓ | — | — | — | — | 0 | — |

### Higher heritability need not imply greater disease relevance

To evaluate the utility of embedding-derived traits with respect to standard uni- and multi-variate traits in a real data setting, we selected the following comparators in the UKB NAFLD cohort:

- Liver fat percentage predictions from a supervised machine learning model trained on neck-to-knee MRIs.
- Embeddings extracted from the penulatimate layer of the above model.
- 203 NAFLD relevant traits with and without adjustment for covariates, treated as a multivariate trait.

Liver fat percentage is known to be a highly predictive biomarker for NAFLD and MRI imputed LF% has previously been used for genetic discovery in NAFLD [15], hence it provides a strong univariate baseline. The 203 NAFLD relevant traits were selected as a non-embedding but multivariate baseline. Many of these traits are expected to be heritable and at least partially disease relevant. To ensure a fair comparison among traits, we only included individuals who had no missing values for any of the traits, thereby ensuring that all GWAS analyses were conducted on the same sample size. For each multivariate trait, we took the first 5 PCs as the input to the EmbedGEM workflow.

Figure 3, shows that while LF% embeddings manifest lower mean *χ*^2^ and fewer genome-wide significant (GWS) hits than the 203 traits, the associations derived from the embeddings have far greater disease relevance. Interestingly, we observe that the LF% embeddings lead to more GWS hits and slightly higher disease relevance than univariate LF predictions, suggesting that embeddings from a performant supervised model might offer more power for genetic discovery than the model’s final predictions. Furthermore, Figure 3 underscores the risks of viewing heritability alone, and in particular the number of GWS associations, as a meaningful gauge of utility for genetic discovery: traits can be highly heritable and yet not disease relevant.

**Fig. 2.**
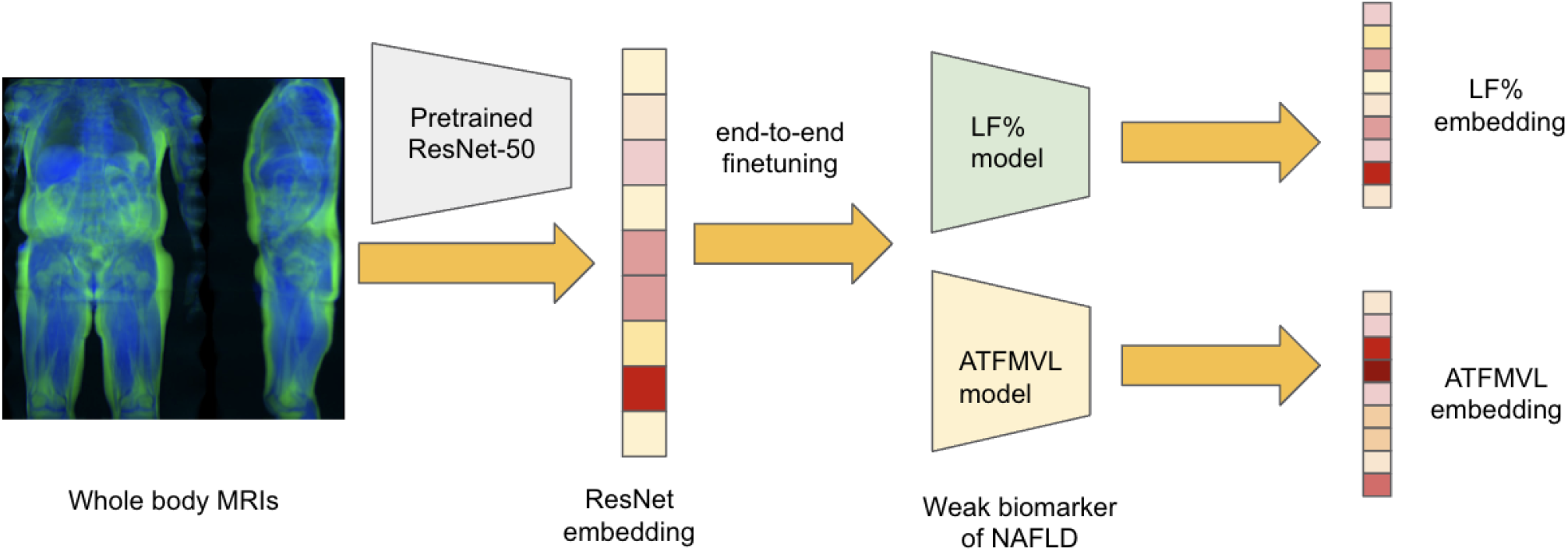
Overview of process used to generate embeddings from neck-to-knee MRIs. An imageNet pre-trained ResNet-50 was used to first generate the ‘ResNet’ embeddings. This model was then subjected to end-to-end finetuning using the method outlined in [15] for two seperate traits, LF% and ATFMVL, to obtain supervised embeddings.

**Fig. 3.**
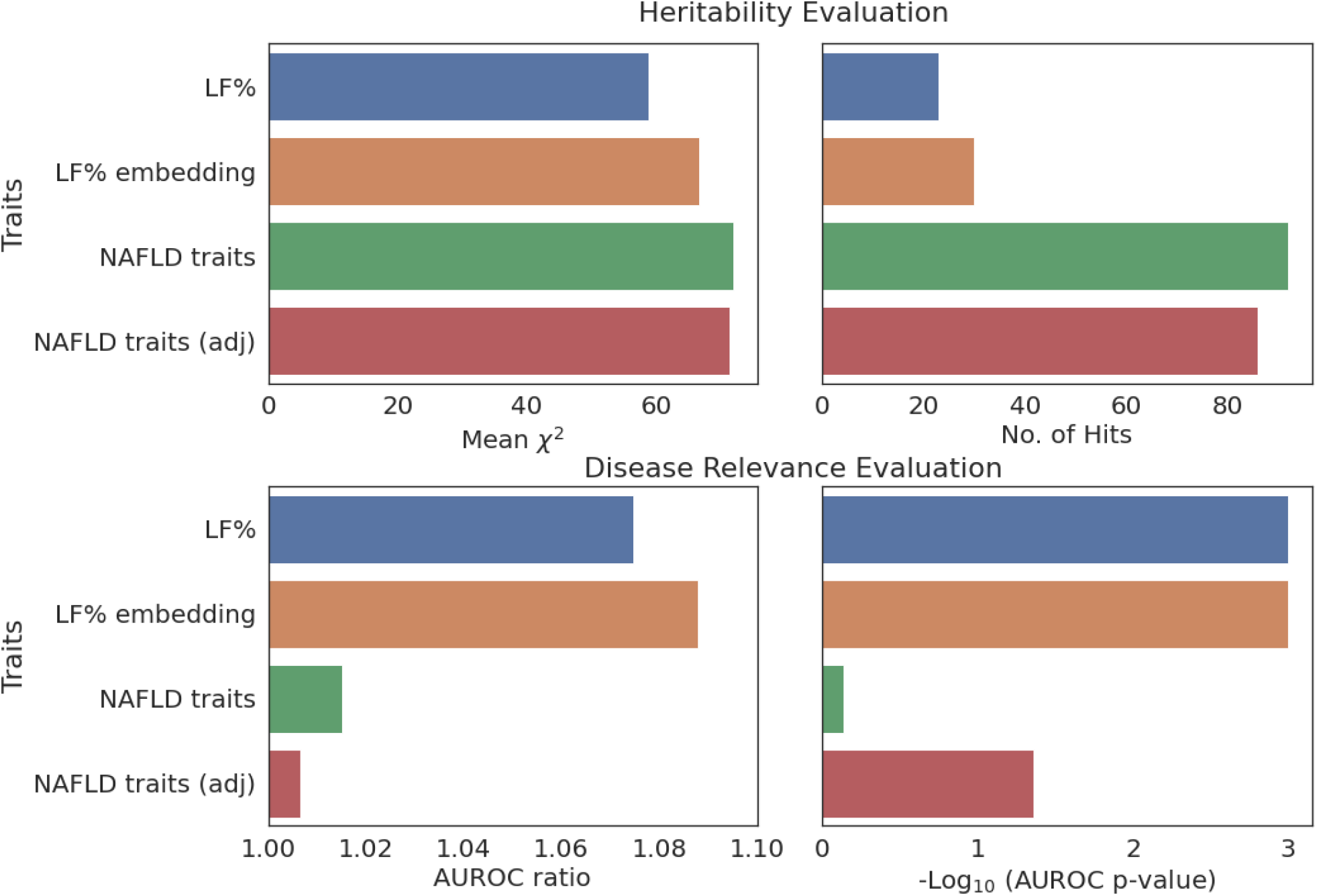
Comparison of heritability and disease relevance between LF embeddings and other traits. The figure illustrates the heritability and disease relevance of liver fat (LF) embeddings, LF percentage predictions, and 203 NAFLD relevant traits. LF embeddings, despite having lower heritability than the 203 traits, show higher disease relevance and heritability than the univariate LF predictions, suggesting that strong supervised embeddings might be more powerful than strong univariate biomarkers.

### Not all embeddings are equally useful

Having demonstrated that, compared to a strong univariate biomarker and non-embedding multivariate traits, embeddings can be both more heritable and more disease relevant, we next examined whether these benefits extend to embeddings trained on less disease-specific tasks. To study this, we compared LF% embeddings with embeddings derived from two other models:

i) a supervised model trained to predict anterior thigh fat-free muscle volume of the left side (ATFMVL) from whole-body MRIs, which has a low correlation with LF% (*r*^2^ = 0.21), and
ii) ImageNet pre-trained ResNet model [10].

As seen in Table 2, embeddings derived from LF% are both more heritable and more relevant to NAFLD than the other embeddings. Note that although 7 variants reached GWS when aggregating across the ImageNet pre-trained embedding PCs, these variants were not GWS for any individual PC, and hence no per-PC PRSs were available for disease relevance evaluation. It seems likely that the paucity of associations with the ImageNet pre-trained embedding is due to the model being tailored for motifs appearing in natural images rather than human biology, or MRIs in particular. Finetuning the ImageNet model on an MRI related task would likely increase embedding heritability, but not necessarily the disease relevance, unless that task was related to NAFLD.

**Table 2.** Comparison of heritability and disease relevance between different embeddings. The table illustrates the heritability and disease relevance of embeddings derived from different machine learning models. We show that depending on how the embeddings are trained, they may have dramatically different utility in genetic discovery. While, a LF% embedding has both high heritability and disease relevance, embeddings of a weaker disease proxy (ATFMVL) has much lower disease relevance and embeddings from ResNet pre-trained on ImageNet show no disease relevance at all.

| Traits | Disease relevance |  |  |  | Heritability |  |
| --- | --- | --- | --- | --- | --- | --- |
| | AUC ratio | AUC p-value | AUPRC ratio | AUPRC p-value | No. of hits | mean $\chi^2$ |
| LF% embedding | 1.088 | 0.001 | 1.162 | 0.001 | 30 | 66.615 |
| ATFMVL embedding | 1.017 | 0.51 | 1.046 | 0.29 | 20 | 33.556 |
| ResNet embedding | — | — | — | — | 7 | 32.440 |

## Conclusion

Here we introduced the first framework specifically intended to evaluate the utility of embeddings for genetic discovery. The genetic validation pipeline is implemented using a combination of several commonly used `plink` commads within a `redun`-based workflow [13]. The workflow comprises two main portions: evaluation of heritability and evaluation of disease relevance. To evaluate heritability, summary statistics from univariate GWAS of orthogonal traits are first aggregated to obtain the equivalent of multivariate GWAS summary statistics, then clumped to obtain independent signals. Metrics for evaluating heritability include the number of independent genome-wide significant (GWS) associations, and the mean *χ*^2^ statistic at GWS loci. To evaluate disease relevance, we examine the collective association of polygenic risk scores (PRSs) for the orthogonalized embeddings with a gold-standard set of labels. Whether the embedding-associated variants significantly improve disease prediction, relative to a set of baseline covariates, is ascertained via a non-parametric paired bootstrapping procedure. The entire workflow can be run end-to-end using a single Python script, available on GitHub at https://github.com/insitro/EmbedGEM, ensuring reproducibility and traceability of results.

We showed that EmbedGEM can successfully disentangle heritability from disease relevance on simulated data. We then demonstrated the utility of EmbedGEM on real data from the UK Biobank using a variety of traits related to non-alcoholic fatty liver disease (NAFLD). We illustrated several important considerations regarding GWAS on embeddings that have not been thoroughly discussed in the literature. First, not all embeddings are equally useful for genetic discovery, and the value of an embedding is tied to the training process by which it was generated. Embeddings adapted to natural images from a ResNet pre-trained on ImageNet show low heritability and (unsurprisingly) no relevance to NAFLD. MRI-adapted embeddings from a model trained to predict an adiposity trait unassociated with NAFLD were more heritable but still not disease relevant. By contrast, embeddings from a MRI-adapted model trained to predict LF%, a key biomarker of NAFLD, were both heritable and significantly disease relevant.

Second, heritability and disease-relevance are separate and distinct properties. A phenotype that yields more genome-wide significant associations is not necessarily more useful for genetic discovery. It is crucial to further examine whether the variants associated with the phenotype are predictive of the trait or disease of ultimate interest. We showed that the heritability of a multivariate collection of 203 traits loosely coupled to NAFLD was significantly higher than that of either ML-derived LF% or LF% embeddings, yet the smaller collection of variants associated with the latter were significantly more disease relevant.

Third, there is evidence that embeddings can in fact facilitate the discovery of disease-relevant genetic variants. Embeddings extracted from a supervised LF% prediction model were more heritable than the original LF% predictions and simultaneously exhibited greater association with NAFLD. A future direction is to develop embeddings tailored for diseases of interest by finetuning foundation models towards the prediction of the disease label itself, or key biomarkers (such as LF %). Our proposed EmbedGEM evaluation framework provides a conceptual and practical means of selecting embeddings that can uncover disease-relevant signals from among alternatives.

Building on this, the software implementation of EmbedGEM has been designed to offer easy extensibility, allowing the computation of additional metrics using the intermediate outputs produced by a highly standardized workflow. For example, users seeking to expand EmbedGEM with novel methods for assessing disease relevance, such as survival analysis or multiple traits, can achieve this by leveraging the PRSs for each PC of the trait. Similarly, if users wish to modify the PRS computation process and employ their own custom tool, they can do so using the clumped summary statistics of each PC. Furthermore, the introduction of new methods for evaluating heritability is feasible by leveraging the multivariate summary statistics generated by EmbedGEM.

We envision EmbedGEM as a key framework for the systematic evaluation of different embeddings and multivariate traits with respect to their utility for genetic discovery. By providing a standardized evaluation workflow, EmbedGEM gives researchers a framework for deciding among various embedding models. We anticipate EmbedGEM will help streamline the process of genetic discovery and encourage reproducibility by enabling results tracking and provenance.

## Acknowledgements

The authors would like thank the participants of the UK Biobank, whose data were used with permission. This research was conducted using the UK Biobank Resource under approved Application Number 51766.

