## Supplementary_materials for "EmbedGEM: A framework to evaluate the utility of embeddings for genetic discovery"

### Supplementary methods

### 1 Genetic data processing and GWAS

To avoid confounding due to population structure, the UK Biobank (UKB) was subset to unrelated subjects of White-British ancestry [9, 3]. Imputed genotypes were filtered to those having a minor allele frequency $> 1\%$, INFO score $> 0.8$, and Hardy-Weinberg equilibrium $P > 1 \times 10^{-10}$. The following standard covariates were included in all GWAS: age, sex, genotyping array, and the top 20 genetic principal components [3]. Genome-wide association studies (GWAS) were performed using `PLINK` (v1.9) [8].

### 2 Config file description

#### 2.1 Heritability module

To evaluate heritability, we start with summary statistics ("sumstats") from an orthogonalized set of traits. This allows users to employ their preferred methods for orthogonalizing the embeddings and performing univariate GWAS. The input sumstats are combined to perform a multivariate test of association, as described in the main paper. The multivariate sumstats are clumped via `plink` with user-specified parameters. After clumping, heritability metrics, including the number of genome-wide significant associations and the mean $\chi^2$ across these associations, are calculated and reported in a `.csv` file. An example of `.yaml` config specifying the parameters relevant to this portion of the workflow is provided below.

```
heritability_eval:
  pval_thresh: 1.0e-3 # default 5e-8
  max_p: 1.0e-1 # default 5e-8

clumping:
  maf_threshold: 0.0001 # default 0.0001
  max_kb: 250 # default 250
  max_pval_all_variants: 0.05 # default 1e-5
  max_pval_lead_variants: 1.0e-3 # default 5e-8
  min_r2: 0.5 # default 0.5
```

The pval_thresh parameter sets the significance threshold and should be ideally set to the same value as the max_pval_lead_variants parameter. The max_p parameter filters out all variants those p-value is higher than this quantity. This can be used to reduce the the size of the multivariate summary statistics file.

#### 2.2 Disease relevance module

To evaluate the disease relevance, sumstats for the orthogonalized embeddings are first clumped, using plink, then combined to form polygenic risk scores (PRSs). Clumping and PRS generation are performed separately for each orthogonalized embedding dimension. These PRSs are then collectively analyzed for association with the gold-standard labels. As for the heritability evaluation, the outputs of this portion of the workflow are reported in a .csv file. Below is an example of specifying the relevant portion of the config .ymal:

```
disease_relevance_options:
  disease_relevance_bfile_template:
  s3://2023-embedgem/toy_data/bfiles/emb_toy_chr{CHROM}
  pheno_file: s3://2023-embedgem/toy_data/test_pheno.tsv
  covar_file: s3://2023-embedgem/toy_data/test_cov.tsv
  pheno_id_col: eid
  cov_id_col: eid
  cov_cols: ['sex']
  pheno_type: continuous
  pheno_col: labels
  pheno_sep: '\t'
  cov_sep: '\t'
  boot_reps: 2000 #default 2000
```

Here, the pheno_type parameter takes the value *binary* or *continuous*, which will determine the type of association test performed and the metrics reported. The boot_reps parameter sets the number of bootstrap replicates for estimating the null distribution of the contrast statistics. The remaining parameters are relevant to the loading of the validation dataset.

### 3 Principal component analysis of liver fat embeddings and NAFLD traits

To motivate retaining only the first several PCs of various multivariate traits, we computed the variance explained by each of the first 20 PCs for the 1024 dimensional LF% embeddings and the 203 NAFLD traits. The results are shown in Figure 1. In each case, more the 80% of the total variance is explained by the first 8 PCs, and the variability contributed by successive PCs rapidly diminishes; the effect is most apparent for the embeddings, where more than half of total variability is explained by the first PC alone.

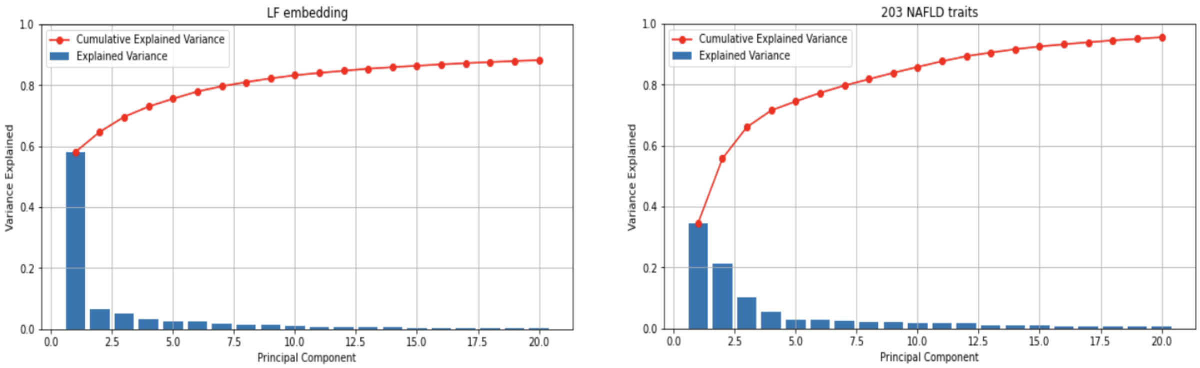

Figure 1: **Rapidly diminishing variability explained by successive principal components.** The figures show the per-component and cumulative variance explained by the first 20 PCs in the case of liver fat percentage embeddings (left) and the collection of 203 NAFLD-relevant traits (right).

### 4 Type 2 diabetes study

#### 4.1 Analysis overview

Type 2 Diabetes (T2D) is a chronic disease characterized by insulin resistance and hyperglycemia, with various resulting complications, including metabolic syndrome, obesity, and optic neuropathy. The global prevalence of T2D is estimated to be 6.1% and increasing, making it a significant public health concern [4].

Machine-learned phenotypes extracted from retinal imaging, including color fundus photography (CFP) and optical coherence tomography, have been associated with various disease outcomes, including age-related macular degeneration, glaucoma, and T2D [6, 7, 1, 5, 11, 10]. In this study, we finetuned the RETFound model [11] to estimate T2D risk from CFPs in the UKB, performed GWAS on the top 10 PCs of the finetuned embeddings, then assessed the utility of the embeddings for genetic discovery via EmbedGEM.

##### 4.2 Model finetuning

RETFound is a foundational vision transformer (ViT) pretrained on $904,170$ CFP images via masked autoencoding [11]. Code for finetuning RETFound is provided with the model release [12]. We finetuned RETFound on $7,820$ CFP images from $3,829$ T2D cases, identified by a E11 ICD10 code, and 7878 images from $3,829$ randomly-selected matched controls, with an HbA1c level below 36 mmol/mol and matching on age and sex. Finetuning was performed on patients of European ancestry, while disease relevance evaluation was performed on patients of non-European ancestry (see below). The train : validation : test split was 55% : 15% : 30%. In the held-out test set, the AUROC and AUPRC for predicting T2D status were 0.68 and 0.67 respectively.

##### 4.3 GWAS

A single image was selected for embedding generation from each of $27,401$ available subjects, following the procedure of Han *et al* [6]. GWAS on the top 10 embedding PCs was performed via linear regression in `plink`. Covariates include age, sex, spherical power, imaging center, image laterality (i.e. right vs left eye), genotyping array, and the the top 20 PCs of the genetic relatedness matrix.

##### 4.4 Evaluation data

The disease relevance evaluation was performed in a cohort of $25,258$ independent patients of non-Europen ancestry who were not included in the GWAS. This cohort included $12,629$

T2D patients, with an E11 ICD10 code, and $12,629$ matched controls, with an HbA1c level below 36 mmol/mol, matching on age and sex.

##### 4.5 Results

After the generation of the T2D embeddings, their evaluation was conducted using the EmbedGEM pipeline. The heritability metrics were as follows:

- Mean $\chi^2$: 80.26
- Median $\chi^2$: 44.42
- Number of hits: 29

The disease relevance metrics were as follows:

- Ratio AUPRC: 0.997 (p-value: $< 0.001$)
- Ratio AUROC: 1.007 (p-value: $< 0.001$)

##### 4.6 EmbedGEM stability anaysis

To evaluate the stability of EmbedGEM metrics to sampling variability in the GWAS data set, we split the discovery cohort into 10 equal folds, staratifying by T2D diagnosis status. Each of the 10 folds was held out in turn, and GWAS was performed on the remaining 9 folds. EmbedGEM was run on the summary statistics from each GWAS. The evaluation data for disease relevance assessment remained the same for all folds. The results are compiled in Table 1. As seen here, the EmbedGEM metrics are quite stable.

#### 5 Computational efficiency of EmbedGEM

The computational efficiency of EmbedGEM was assessed across nested variant sets of increasing size (1K, 10K, 100K variants), each subset from the pool of common variants within the UKB. The size of the GWAS cohort was fixed at 10K, and evaluation cohort sizes of 1K and 10K were examined. The bootstrap resampling procedure was iterated 100 times for

| Excluded fold | Ratio AUPRC | Ratio AUROC | Mean Chi2 | Median Chi2 | No. of Hits |
|---|---:|---:|---:|---:|---:|
| Fold 0 | 1.003 | 1.028 | 74.1 | 41.2 | 29 |
| Fold 1 | 1.005 | 0.999 | 81.7 | 45.5 | 25 |
| Fold 2 | 1.005 | 1.000 | 76.8 | 47.0 | 29 |
| Fold 3 | 0.997 | 1.010 | 83.7 | 55.0 | 26 |
| Fold 4 | 0.999 | 1.008 | 77.9 | 47.7 | 23 |
| Fold 5 | 1.002 | 1.001 | 79.8 | 49.5 | 24 |
| Fold 6 | 1.006 | 1.002 | 78.3 | 38.5 | 31 |
| Fold 7 | 0.998 | 1.008 | 76.1 | 43.4 | 26 |
| Fold 8 | 0.998 | 1.007 | 74.1 | 36.8 | 29 |
| Fold 9 | 1.000 | 1.002 | 80.3 | 44.4 | 29 |
| Mean (SE) | 1.001 (0.001) | 1.007 (0.003) | 78.3 (1.0) | 44.9 (1.7) | 27.1 (0.8) |

Table 1: **Evaluating stability of EmbedGEM metrics**: We split the discovery cohort in T2D into 10 folds and used 9 of the folds to perform GWAS and generate EmbedGEM metrics. We report the mean and standard error of each metrics at the bottom.

each analysis. All computations were performed on a p3.2xlarge instance on the Amazon Web Services (AWS) Elastic Compute Cloud (EC2). End-to-end run times are presented in Table 2. EmbedGEM exhibited no marked increase in computational time either with increasing numbers of variants or increasing numbers of subjects in the evaluation data set. Further investigation suggests that the end-to-end run times are dominated by the overhead associated with multi-threading and launching batch jobs, rather than the computational complexity of the EmbedGEM analysis.

| | **1K SNPs** | **10K SNPs** | **100K SNPs** |
|---|---|---|---|
| **1K Individuals** | 192.98 s | 193.02 s | 194.01 s |
| **10K Individuals** | 191.93 s | 189.71 s | 192.14 s |

Table 2: **End-to-end runtimes of EmbedGEM across different numbers of variants and different evaluation cohort sizes.**

To elucidate the computational demands of individual EmbedGEM components, we quantifie the run times across varying input dimensions, as depicted in Figure 2. Size, as depicted on the X-axis, refers generically to the dimension along which the run time scaled most quickly. For GWAS, this was the number of subjects. Based on the slope in Figure 2, a 10× increase in the sample size resulted in a 7.8× increase in run time. For clumping and PRS calculation, the scaling dimension was the number of variants. 10× increases were associated with 3.3×

and 6.3× fold increases in run time, for clumping and PRS respectively. For calculating disease relevance metrics, the scaling dimension was the number of bootstraps. A 10× increase was associated with a 4.4× increase in run time. Note that for disease relevance, run times were evaluated at smaller sizes because the run time was relatively large even at only 100 bootstraps.

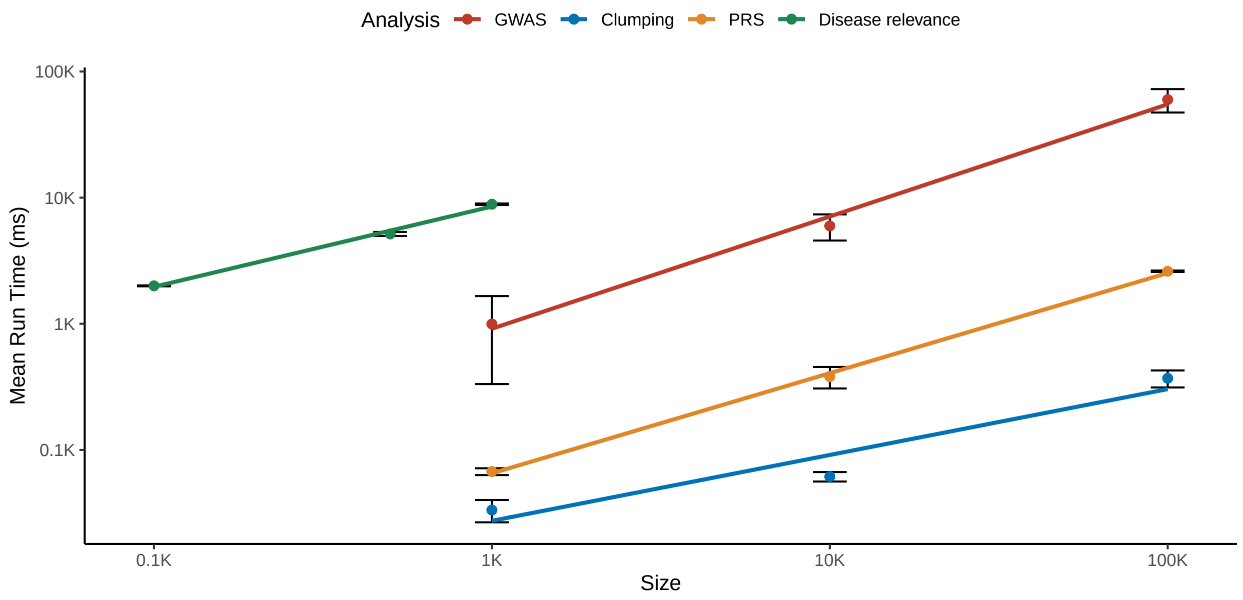

Figure 2: **Run times for specific components of the EmbedGEM evaluation.** Size refers generally to the dimension along which the run time of the analysis scales most quickly. For GWAS, this is the number of subjects. For clumping and polygenic-risk score (PRS) calculation, this is the number of variants. For the disease relevance evaluation, this is the number of bootstrap samples.

### 6 Proxies for heritability

#### 6.1 Expected $\chi^2$ statistic and number of significant hits

For a univariate, quantitative trait $Y$, the standard heritability model partitions variation in $Y$ into two sources, genetic $\gamma$ and environmental $\epsilon$:

$$Y = \gamma + \epsilon.$$

The broad sense heritability is defined as the proportion total phenotypic variation that is due to genetic variation:

$$H^2 = \frac{\mathbb{V}(\gamma)}{\mathbb{V}(Y)}$$

When estimating SNP heritability, as in common in GWAS, the genetic component is decomposed as the sum of many individual variants with infinitesimal effects:

$$\gamma = \sum_{j=1}^{M} G_j \beta_j.$$

Here $G_j$ is the number of minor alleles at the $j$th variant, $M$ is the total number of causal variants, and the effect sizes $\beta_j$ are assumed to follow a normal distribution with mean 0 and per-SNP variance $\sigma^2_{\text{SNP}}$:

$$\beta_j \overset{\text{IID}}{\sim} N(0, \sigma^2_{\text{SNP}}). \tag{1}$$

Assuming further that *i.* the causal variants are in linkage equilibrium, that *ii.* the effect sizes are independent of the minor allele frequency, and that *iii.* the variants are standardized to have mean 0 and variance 1, the genetic variance reduces to:

$$\mathbb{V}(\gamma) = \mathbb{V}\left(\sum_{j=1}^{M} G_j \beta_j\right) \overset{i.}{=} \sum_{j=1}^{M} \mathbb{V}(G_j \beta_j) \overset{ii.}{=} \sum_{j=1}^{M} \mathbb{V}(G_j)\mathbb{V}(\beta_j) \overset{iii.}{=} M \times \sigma^2_{\text{SNP}},$$

where the Roman numeral above the equality shows how each assumption was used. Overall, under the proposed model, the SNP heritability is:

$$h^2 = \frac{\mathbb{V}(\gamma)}{\mathbb{V}(Y)} = \frac{M\sigma^2_{\text{SNP}}}{\mathbb{V}(Y)}.$$

Thus, we see already that the heritability of a trait is directly proportional to the number of causal variants $M$. Now, for the $j$th genetic variant, the $\chi^2$ statistic for association with the phenotype is:

$$\chi_j^2 = \left\{ \frac{\hat{\beta}_j}{\mathbb{SE}(\hat{\beta}_j)} \right\}^2$$

where $\hat{\beta}_j$ is the estimated effect and $\mathbb{SE}(\hat{\beta}_j)$ is its standard error (SE). Since the SE of the effect size is inversely proportional to the square root of the sample size:

$$\mathbb{SE}(\hat{\beta}_j) \propto \frac{1}{\sqrt{N}},$$

the $\chi^2$ statistic is directly proportional to the sample size and the square of the effect size:

$$\chi_j^2 \propto \frac{\hat{\beta}_j^2}{1/N} = N \times \hat{\beta}_j^2.$$

Under the infinitesimal model in (1):

$$\mathbb{E}(\hat{\beta}_j^2) = \sigma^2_{\text{SNP}}. \tag{2}$$

In the heritability module, we calculate the mean $\chi^2$ statistic, denoted $\overline{\chi^2}$, by taking the mean of the per-variant $\chi^2$ statistics *after* clumping to remove linkage disequilibrium:

$$\overline{\chi^2} = \frac{1}{M_{\text{clump}}} \sum_{j=1}^{M_{\text{clump}}} \chi_j^2.$$

With (2), the expectation of the mean $\chi^2$ statistic is:

$$\mathbb{E}(\overline{\chi^2}) = \frac{1}{M_{\text{clump}}} \sum_{j=1}^{M_{\text{clump}}} \mathbb{E}(\chi_j^2) = \mathbb{E}(\chi_j^2) \propto \sigma^2_{\text{SNP}} \times N$$

Since the SNP heritability $h^2$ and the mean $\chi^2$ are each proportional to $\sigma^2_{\text{SNP}}$, it follows that:

$$h^2 = \frac{M\sigma^2_{\text{SNP}}}{\mathbb{V}(Y)} \propto \frac{M}{\mathbb{V}(Y)} \frac{\mathbb{E}(\overline{\chi^2})}{N}. \tag{3}$$

Thus, the SNP heritability is directly proportional to the expected $\chi^2$ statistic. There are several sources of error in this analysis. First, as is common when estimating SNP heritability, we have ignored both gene-environment interaction and non-additive sources of genetic variation, including dominance and epistasis. Second, we have assumed that causal variants for a trait are in linkage equilibrium. This assumption is immaterial insofar as the more careful analysis by Bulik-Sullivan *et al* [2], which accounts for linkage disequilibrium (LD) among causal variants, reaches the same conclusion. In fact, the direct proportionality between the expected $\chi^2$ statistic and $h^2$ is the basis for heritability estimation via LD score regression [2]. Third, the number of variants that remain after clumping $M_{\text{clump}}$ is expected to provide only a lower bound of the true number of causal variants $M$ since, in finite samples, not all causal variants will reach genome-wide significance. Despite these limitations, the number of genome-wide significant associations (after clumping) and the expected $\chi^2$ statistic remain useful proxies for SNP heritability due to their ease of calculation and direct proportionality with $h^2$.

#### 6.2 Empirical comparison of the relationship between heritability and our proxy measures

We selected a subset of 10 relatively uncorrelated biomarker and metabolite traits, from among the 203 NAFLD-relevant traits described in the main text, and performed GWAS, clumping, and mean $\chi^2$ estimation. We then estimated the SNP heritability of the these same phenotypes via LD score regression [2], and compared it with our proxy metrics. Consistent with the relationship in equation (3), we observe a relatively strong linear association between LD score regression's estimate of heritability and our proxy metrics.

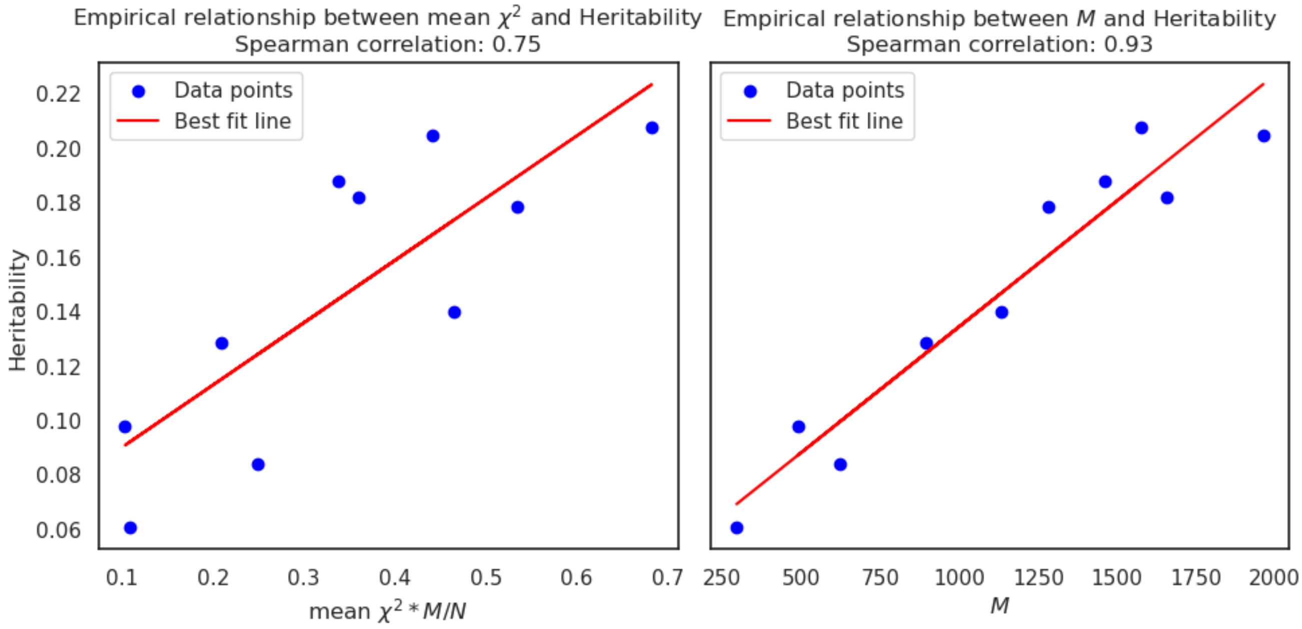

Figure 3: **Comparing the theoretical estimates of heritability using mean $\chi^2$ and $M$ with heritability estimates from LD score regression [2].**
